## Supplementary information for "Taste of time: A porous-medium model for human tongue surface with implications for early taste perception"

**Supplementary Tables**

**Supplementary Table S1.** Intensity perception ratings and simulated stimulus concentrations for different pulse durations of NaCl and NaSac stimuli

| **Measure** | **500mM NaCl solution** | | | | **2mM NaSac solution** | | | |
| --- | --- | --- | --- | --- | --- | --- | --- | --- |
| Intended pulse duration (ms) | 100 | 200 | 300 | 1000 | 100 | 200 | 300 | 1000 |
| Intensity rating^a^ | 11 | 14 | 12 | 18 | 8 | 12.5 | 10.5 | 16 |
| Simulated tongue concentration (mM)^b^ | 211.81 | 416.61 | 460.05 | 494.48 | 0.85 | 1.67 | 1.84 | 1.98 |

^a^ Data were from Kelling and Halpern’s intensity rating experiment, Ref.^1^ A 2s stimulus pulse was assigned a modulus of 20 and used as a standard against which subjects would judge in proportion of the intensity of the experimental stimulus.

^b^ The correlation between the simulated tongue concentration and the experimental intensity ratings for NaCl and NaSac stimulus pulses were 0.70 and 0.82, respectively

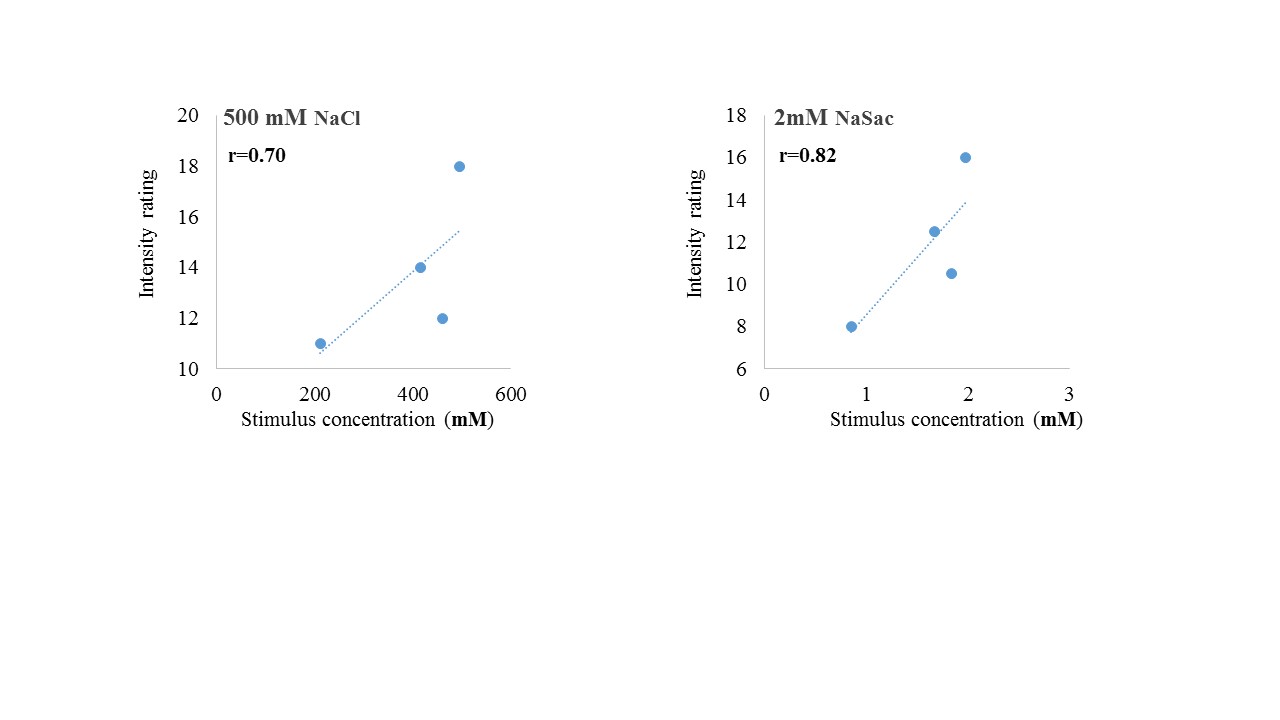

**Supplementary Table S2.** Peaks of intensity perceptions and simulated peak stimulus concentrations in the tongue surface zone for various setups

| Measure | Sucrose solution^a^ | | | | NaCl solution^b^ | | | | | |
| --- | --- | --- | --- | --- | --- | --- | --- | --- | --- | --- |
| Stimulus solution concentration (mM) | 100 | 180 | 320 | 560 | 150 | 300 | 450 | 150 | 300 | 450 |
| Holding duration (s) | 10 | 10 | 10 | 10 | 5 | 5 | 5 | 10 | 10 | 10 |
| Normalized Peak intensity rating | 2.1 | 3.5 | 6.1 | 6.9 | 20.2 | 30.0 | 41.8 | 22.5 | 35.8 | 51.7 |
| Simulated tongue concentration (mM)^c^ | 20.7 | 37.3 | 66.2 | 115.9 | 38.8 | 77.6 | 116.5 | 54.8 | 109.5 | 164.3 |

^a^ The sucrose solution intensity ratings were abstracted from the median ration judgements of taste intensity test by Lawless and Skinner, Ref.^2^. The subjects in the intensity rating test were taught to move a pointer which could mark off a distance from zero proportional to the taste intensity. They were told to estimate ratios, so that if the taste intensity doubled from some level, the pointer would be moved twice as far from zero. Similarly, the subjects were instructed to preserve ratio properties between stimuli, such that if the second stimulus was half as strong at its peak as the first, the pointer would be moved half as far from zero.

^b^ The NaCl solution intensity ratings were from Matuszewska et al.’s work, Ref.^3^ Two reference standards were evaluated (distilled water and 2.5% NaCl solution) before each experiment in order to determine the range of intensity scale, which is from ‘none, 0’ to ‘very intensive, 100’.

^c^ The correlation between the simulated tongue concentration and the experimental peak intensity ratings for sucrose solution and NaCl solution were 0.933 and 0.993, respectively.

**Supplementary Table S3.** Estimated diffusivity of selected sweeteners and mean time-intensity parameters

| Measure | Saccharin | Sucrose | Aspartame | Acesulfame K | Cyclamate | Sucralose | Crystalline  fructose |
| --- | --- | --- | --- | --- | --- | --- | --- |
| Diffusivity  (×10^–10^ m^2^/s)^a^ | 7.06 | 4.76 | 4.64 | 7.73 | 6.62 | 4.55 | 7.00 |
| T_max_^b^ (s) | 2.72 | 3.22 | 3.02 | 2.15 | 2.97 | 2.88 | 2.95 |
| M_abs_^c^ | 10.46 | 10.39 | 7.11 | 12.95 | 11.74 | 9.72 | 11.66 |
| T_lag_^d^ (s) | 0.78 | 0.63 | 0.92 | 0.55 | 0.72 | 0.82 | 0.72 |

^a^ All the diffusivities were estimated using Wilke and Chang equation, Ref.^4,5^.

^b^ Time to reach peak perceived intensity, Ref.^6^ The correlation coefficient between T_max_ and diffusivity is –0.689 (*p* = 0.087).

^c^ Slope to reach peak intensity, Ref.^6^ The correlation coefficient between M_abs_ and diffusivity is 0.789 (*p* = 0.035).

^d^ Lag time between ingestion and onset of response, Ref.^6^ The correlation coefficient between T_lag_ and diffusivity is -0.545 (*p* = 0.206).

**Supplementary Figure**

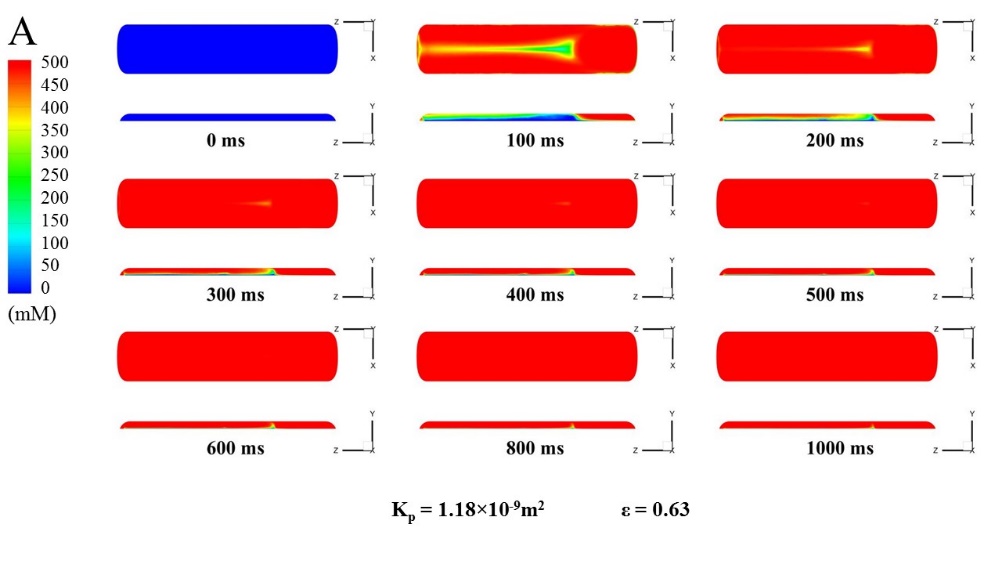

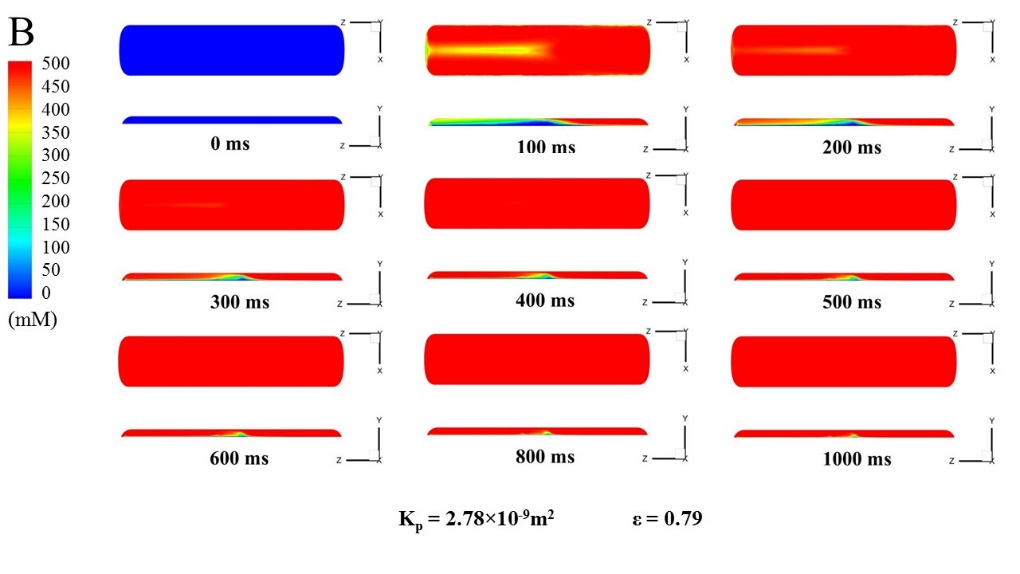

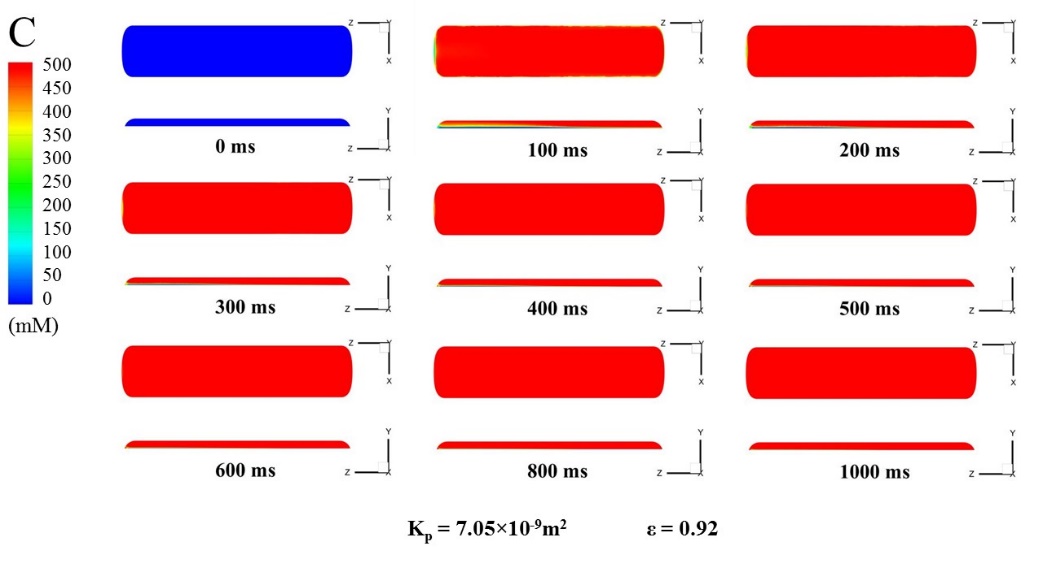

**Supplementary Figure S1.** Top-surface view (top row) and cross-section view (bottom) of simulated NaCl concentration profile throughout the papilla layer at different time points, with stimulus of 500mM NaCl: **(A)** case 1: K_p_ (permeability)=1.18×10^-9^ m^2^, ε (porosity)=0.63; **(B)** case 2: K_p_=2.78×10^-9^ m^2^, ε=0.79; **(C)** case 3: K_p_=7.05×10^-9^ m^2^, ε=0.92. Parameters for each case are listed in Table 1.
